# BGC Atlas: A Web Resource for Exploring the Global Chemical Diversity Encoded in Bacterial Genomes

**DOI:** 10.1101/2024.08.23.609335

**Authors:** Caner Bağcı, Matin Nuhamunada, Hemant Goyat, Casimir Ladanyi, Ludek Sehnal, Kai Blin, Satria A. Kautsar, Azat Tagirdzhanov, Alexey Gurevich, Shrikant Mantri, Christian von Mering, Daniel Udwary, Marnix H. Medema, Tilmann Weber, Nadine Ziemert

## Abstract

Secondary metabolites are compounds not essential for an organism’s development, but provide significant ecological and physiological benefits. These compounds have applications in medicine, biotechnology, and agriculture. Their production is encoded in biosynthetic gene clusters (BGCs), groups of genes collectively directing their biosynthesis. The advent of metagenomics has allowed researchers to study BGCs directly from environmental samples, identifying numerous previously unknown BGCs encoding unprecedented chemistry. Here, we present the BGC Atlas (https://bgc-atlas.cs.uni-tuebingen.de), a web resource that facilitates the exploration and analysis of BGC diversity in metagenomes. The BGC Atlas identifies and clusters BGCs from publicly available datasets, offering a centralized database and a web interface for metadata-aware exploration of BGCs and gene cluster families (GCFs). We analyzed over 35,000 datasets from MGnify, identifying nearly 1.8 million BGCs, which were clustered into GCFs. The analysis showed that ribosomally synthesized and post-translationally modified peptides (RiPPs) are the most abundant compound class, with most GCFs exhibiting high environmental specificity. We believe that our tool will enable researchers to easily explore and analyze the BGC diversity in environmental samples, significantly enhancing our understanding of bacterial secondary metabolites, and promote the identification of ecological and evolutionary factors shaping the biosynthetic potential of microbial communities.

## 1 Introduction

Secondary (or specialized) metabolites are biomolecules that are not essential for the growth or survival of an organism, but provide important ecological and physiological advantages to it. Secondary metabolites produced by bacteria have a wide array of functions, including acting as antibiotics, antifungals, anticancer agents, and immunosuppressants [1]. They are also important in agriculture as plant growth promoters [2] and in the biotechnology industry for various applications [3]. The ecological roles of secondary metabolites include mediating interactions within microbial communities, such as communication between microbes or between microbes and their hosts [4].

The production of secondary metabolites is controlled by biosynthetic gene clusters (BGCs), which are sets of genes closely located in a genome [5]. These clusters coordinate the production of secondary metabolites by encoding the necessary enzymes and regulatory proteins. BGCs are often complex, and include not only the genes for the biosynthetic pathway, but also regulatory elements and sometimes resistance genes to protect the organism from the toxic effects of its own products [6, 7].

Historically, the study of secondary metabolites and their biosynthetic pathways has relied on the cultivation of microorganisms in the laboratory. However, this approach has significant limitations. Many microorganisms are not easily cultured under laboratory conditions [8], and even those that can be cultured may not express their full biosynthetic potential in artificial environments [9]. This has led to a significant underestimation of the diversity of secondary metabolites in nature.

The advent and widespread adoption of metagenomics have revolutionized the study of BGCs and their associated secondary metabolites, enabling direct genetic analysis from environmental samples without the need for culturing [10, 11]. This approach has uncovered a vast array of previously unknown BGCs across diverse environments, such as soil [12], marine ecosystems [13], and the human microbiome [14], revealing significant potential for discovering novel secondary metabolites with important biological and industrial applications [15]. Through metagenome mining approaches, successful attempts have also been made to recover culture-independent natural products from environmental samples. For example, malicidines, a class of antibiotics active against multidrug-resistant pathogens [16], and functionally and structurally diverse type II polyketide synthase enzymes have been discovered from environmental soil samples [17].

Despite these advances, the exploration and analysis of BGCs in metagenomic datasets remain challenging. The complexity and sheer volume of metagenomic data require sophisticated bioinformatic tools and resources to identify, annotate, and analyze BGCs. Furthermore, integrating BGC data with environmental metadata is crucial for understanding the ecological and evolutionary contexts of secondary metabolite production.

Although there exist other resources which gather BGCs from human microbiome samples, or publicly available isolate genomes or metagenome-assembled genomes (MAG), they do not focus on the analysis of environmental samples from a wide range of environments as a whole, and their association with environmental metadata. ABC-HuMi [18] focuses on human microbiomes, while antiSMASH-DB [19] and BiG-FAM [20] contain data from cultured microbial genomes or MAGs deposited in public repositories. Additionally, sBGC-hm [21] is specifically dedicated to MAGs from human gut microbiomes

To address these challenges, we developed the BGC Atlas, a comprehensive web resource for exploring the diversity of biosynthetic gene clusters in metagenomes with corresponding environmental metadata. The BGC Atlas leverages publicly available metagenomic sequencing datasets and integrates advanced computational tools for BGC identification and clustering. This resource provides a centralized database for exploring BGCs, gene cluster families (GCFs), and their associations with environmental sample metadata, enabling researchers to explore BGC diversity in the context of the environments from which they are derived. Users can query the database for similarities to identified BGCs, analyze the diversity of GCFs in specific environments, and explore distribution patterns of BGCs of interest in relation to environmental factors. The BGC Atlas offers a significant advancement in the study of bacterial secondary metabolites, offering researchers a powerful tool to explore and analyze the biosynthetic potential of microbial communities with reference to their natural environments. It aims to serve as a comprehensive resource to enhance our understanding of secondary metabolites and their impact on ecological and evolutionary processes, and map the chemical diversity of the world encoded in bacterial genomes.

## 2 DATA COLLECTION, PROCESSING, AND ANALYSIS

### 2.1 Data Collection and Preprocessing

The initial step in our workflow involved the collection of metagenomic datasets and their corresponding metadata from public repositories. We sourced our data from MGnify [22], a comprehensive database providing access to tens of thousands of metagenomic samples across various environments. The datasets collected encompassed diverse habitats, including marine, soil, human-associated, and other environmental samples.

For each metagenomic sample in the MGnify database, we collect assembled contigs in FASTA format, as well as associated metadata that provides details describing the environmental and biological context of the samples from which these metagenome assemblies are produced (Figure 1).

**Figure 1.**
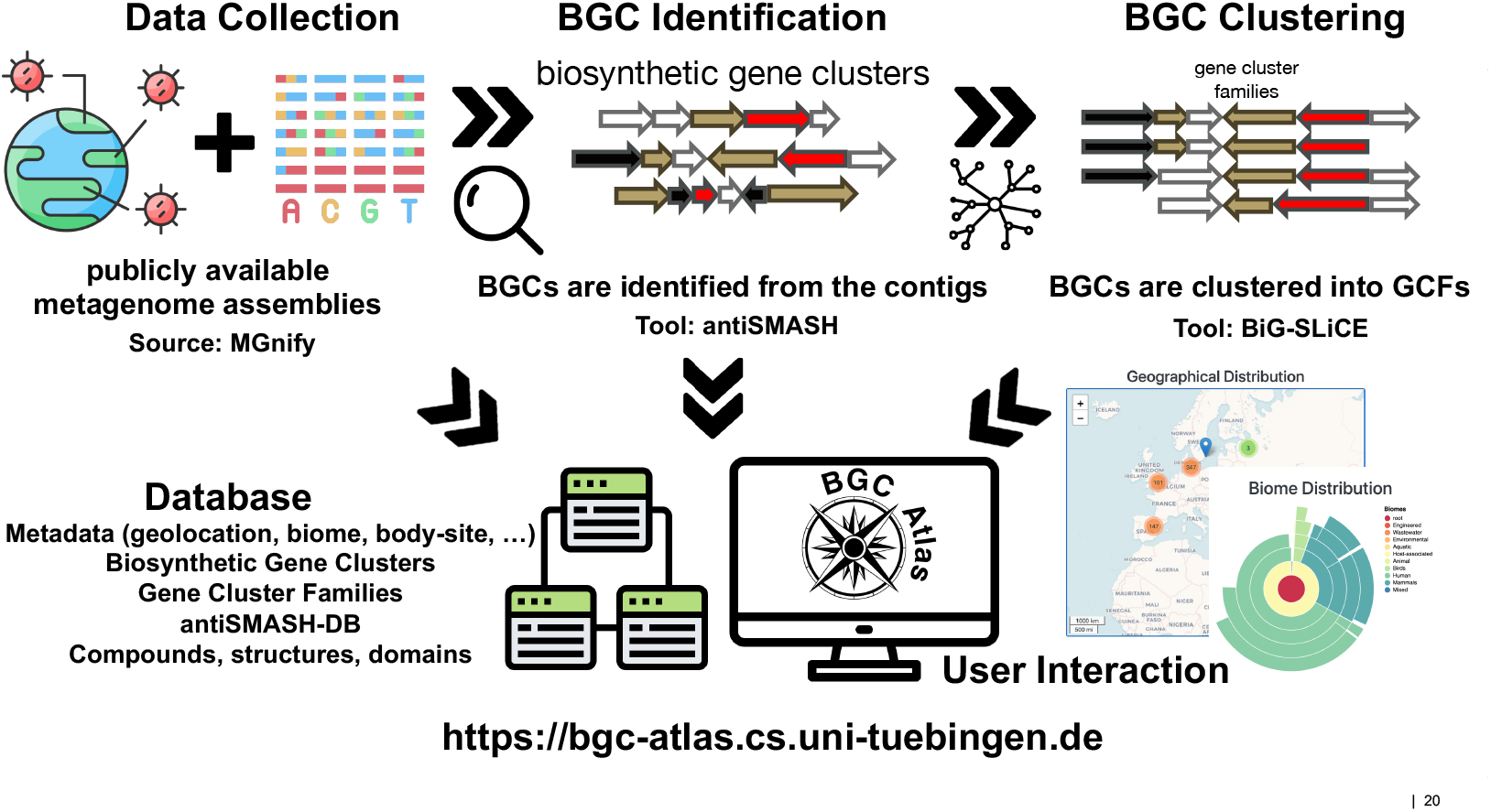
The BGC Atlas data processing pipeline. The BGC Atlas was developed as a comprehensive web resource by using publicly available metagenomic datasets from MGnify. The process involved collecting metagenomic assemblies and metadata, identifying biosynthetic gene clusters (BGCs) using antiSMASH, clustering these BGCs into gene cluster families (GCFs) with BiG-SLiCE, and storing the results in a PostgreSQL database. A userfriendly web interface was then created to assist data exploration, and to query for similar BGCs.

### 2.2 BGC Identification

The identification of biosynthetic gene clusters (BGCs) was carried out using antiSMASH [23] (version 7.0.0, with parameters `--clusterhmmer --tigrfam --asf --cc-mibig --cb-subclusters --cb-knownclusters --pfam2go --rre --tfbs --genefinding-tool prodigal-m --allow-long-headers` as described in [19]). antiSMASH with this set of parameters provides comprehensive annotations, including the type of secondary metabolite produced, predicted chemical structures, and functional domains present within the BGCs, and makes it possible to build an antismash-DB [19] for the predicted BGCs, which organizes and stores detailed information about the predicted BGCs, enabling further comparative analyses and data retrieval (Figure 1).

### 2.3 BGC Clustering

To manage the large number of identified BGCs and to gain meaningful biological insights, we clustered the BGCs into gene cluster families (GCFs) using BiG-SLiCE (version 2.0.0) [24]. BiG-SLiCE is a scalable tool that groups BGCs based on their sequence similarity and domain architecture, allowing for the formation of non-redundant sets of GCFs. These GCFs represent groups of BGCs that are predicted to produce structurally similar secondary metabolites (Figure 1).

The clustering process involved two main steps:

1. **Initial Clustering:** We first clustered BGCs annotated as complete by antiSMASH. This step ensured that the core clusters were well-defined and represented the primary biosynthetic capabilities within the datasets. The initial clustering was performed using a distance threshold of 0.4, a value chosen to balance the grouping of similar BGCs while avoiding excessive clustering that could obscure biological relevance.
2. **Assignment of Partial BGCs:** In a secondary step, partial BGCs (annotated as “on contig edge” by antiSMASH) and known compounds from the MIBiG database [25] were assigned to the predefined clusters from the first step, using the BiG-SLiCE query mode. This enhanced the comprehensiveness of our GCFs, incorporating a broader range of BGCs and ensuring a more complete representation of biosynthetic diversity. In this step, we used a distance threshold of 0.4, as in the initial clustering, and retained only the best hit in case a BGC was assigned to multiple GCFs. We have also retained all those hits above the threshold of 0.4, and report them as putative together with their distances.

### 2.4 Database and Web Interface

The results from the BGC identification and clustering processes are stored in a PostgreSQL database. This relational database system was chosen for its robustness, scalability, and ability to handle complex queries efficiently. The database schema was designed to accommodate the diverse types of data generated, including BGC sequences, annotations, clustering information, and associated environmental metadata (Supplementary Figure 1). For users requiring programmatic access, we provide a complete database dump for download on our website.

To provide researchers with easy access to the data, we developed a web interface for the BGC Atlas (Figure 2). It was built using modern web development frameworks (node.js, express, pug, leaflet, and bootstrap), ensuring a responsive and user-friendly experience. Key features of the web interface include:

**Figure 2.**
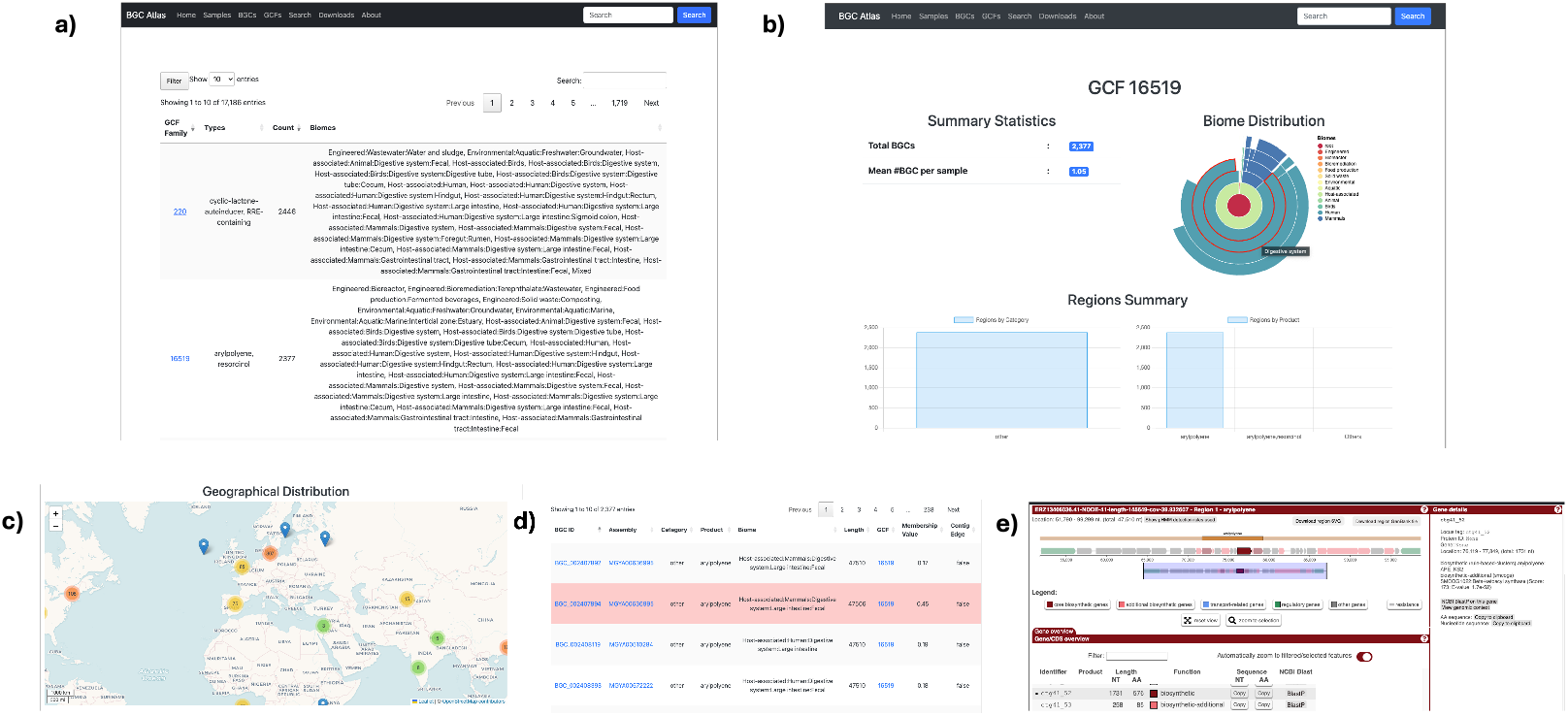
Example visualization from the BGC Atlas web interface. a) The table view for gene cluster families. b) The BGC view, filtered for a specific GCF, shows summary statistics and biome distribution, as well as the product types for the selected GCF. c) The geographical distribution of samples in which the filtered GCF can be found. d) The list of BGCs belonging to this GCF, links to their sample and individual BGC pages, and their associated metadata. e) The antiSMASH view of a selected BGC.

- **BGC and GCF Exploration:** Users can browse the identified BGCs, view detailed annotations, and explore their distribution across different environments.
- **Metadata Integration:** The interface allows users to connect BGC data with environmental metadata to enable ecological and evolutionary analyses and hypothesis generation.
- **Query Functionality:** Users can input their own BGC sequences to search for similar clusters within the database. This feature uses BiG-SLiCE’s query mode, which compares the input BGCs to those in the database based on domain architecture and sequence similarity.

The results of the comparison are presented in a table, showing the distance from the input BGC’s to its closest matching GCF in the database.

The BGC Atlas provides several views and functionalities to explore metagenomic samples, BGCs, and GCFs. All of these views are searchable and filterable for either free text or available metadata features. The “Filter” menu can combine the boolean logic of multiple expressions, such as searching for “Marine” (biome feature) BGCs from “Arctic” (geolocation feature) environments. The “Samples” view provides an overview of all analyzed samples with their metadata features, and lets users to explore them within the antiSMASH interface. The “BGCs” and “GCFs” views display tables of all BGCs or GCFs stored in the database, with filtering options available for more specific queries. In the “GCFs” view, selecting a specific GCF filters the “BGCs” view to display only its member BGCs. This lets users to transition between views and examine the detailed annotations and distributions of BGCs within the context of their respective GCFs, allowing them to perform in-depth comparative analyses and explore metadata to point at ecological roles (Figure 2).

## 3 Results

### 3.1 Preliminary Data Insights

#### 3.1.1 Distribution of BGC Structural Classes

Our initial analysis of 35,486 datasets from MGnify revealed the presence of approximately 1.85 million BGCs across diverse environmental samples. A vast majority (88.7%) of the identified BGCs are annotated as “on contig edge”, by antiSMASH, suggesting their incompleteness, due to the nature of metagenomic sequencing which often results in short contigs. However, more than half of the BGCs that we identified (51.7%) are still longer than 5kbp, with those that are complete having a median length of approximately 21kbp.

The majority of the BGCs identified in the datasets that we analyzed (52.3%) were ribosomally synthesized and post-translationally modified peptides (RiPPs), with cyclic-lactone-autoinducer being the most common product type (16.5%). Other notable structural classes include terpenes, non-ribosomal peptides, and polyketides, contributing to the chemical diversity observed in the analyzed metagenomes. We also observed distinct distribution patterns of product categories across different biomes. For instance, terpenes occur predominantly in terrestrial environmental samples, while RiPPs emerge as the most abundant class in host-associated samples (Supplementary Figure 2).

#### 3.1.2 Habitat Specificity of Gene Cluster Families

Our clustering analysis revealed that GCFs exhibit significant habitat specificity, indicating that certain BGCs are adapted to specific environmental conditions. Overall, 76% of GCFs with at least five BGC members were found exclusively in one specific habitat, emphasizing the ecological specialization of microbial secondary metabolism (Figure 3).

**Figure 3.**
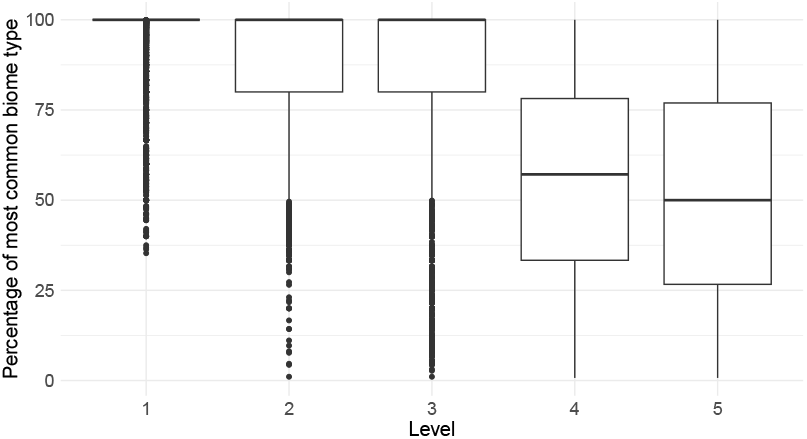
: Box plots illustrating the proportion of the most common biome type for each GCF with at least five members across different hierarchical levels of biome ontology. Each level represents a distinct stage in the hierarchical classification of biomes (e.g. Level 1: Host-associated, Level 2: Humans, Mammals, and Birds). The figure highlights that GCFs are highly specific to the biomes that they appear in, especially at higher levels of the biome ontology.

We observe that a vast majority of GCFs are extremely specific to their habitat, especially at the higher levels of the biome ontology provided by MGnify. At the first level of the hierarchy (Host-associated, Environmental, Engineered, Mixed), almost all GCFs are found exclusively in one of the categories (Figure 3). This trend still holds for a significant majority of GCFs at the second (e.g. Human vs. Mammals for Host-associated) and the third (e.g. Digestive system vs. Skin for Human) levels, and starts to decrease as the habitat becomes more specialized (e.g. Large intestine vs Oral for Human-Digestive system at level 4).

Detailed examination of certain GCFs also revealed some remarkable patterns of habitat specificity. For example, GCF5261, a terpene-producing cluster, was found exclusively in river estuaries across the globe, highlighting its adaptation to this specific environment. On the other hand, GCF12475, a family of NRPS BGCs approximately 50kb in length, was identified solely in mammalian faecal samples from Europe, Southeast Asia, and North America. The ecological roles of these specialized BGCs suggest potential adaptive strategies employed by microbes to thrive in their respective environments.

## 4 Discussion and Future Directions

The BGC Atlas significantly advances the study of secondary metabolites, by offering a comprehensive resource for the exploration and analysis of BGCs in metagenomes. By integrating large-scale metagenomic data and their meta-data with state-of-the-art bioinformatics methods, the BGC Atlas provides a novel view of the diversity and distribution of BGCs across different environments.

Secondary metabolites have significant ecological implications. These compounds play crucial roles in microbial ecology, mediating interactions within microbial communities, and between microbes and their hosts. Our initial findings suggest that microbes may employ specific BGCs as adaptive strategies to thrive in distinct ecological niches. The habitat specificity observed in many gene cluster families underscores the potential for secondary metabolites to influence microbial community structure, faciliate interspecies communication, and contribute to overall ecosystem stability. Furthermore, this specificity hints at the evolutionary pressures that drive the diversification of BGCs, potentially leading to novel bioactive compounds with unique biological activities.

While the BGC Atlas offers significant advancements in bacterial secondary metabolite research, it is also important to note its certain limitations. It is currently limited to the datasets in the MGnify database. In the future, we plan to expand our data resources by incorporating more metagenomic datasets from additional publicly available resources such as JGI IMG/M [26], Logan [27], as well as through collaborative sequencing efforts, to enhance the depth of the BGC Atlas. Currently, most of our analysis is limited to short-read sequencing datasets, often resulting in BGCs being located at contig edges, complicating the accurate reconstruction and annotation of these clusters. We expect with the growing amount of long-read sequencing data in the repositories, this issue will be mitigated, with longer contigs containing more complete BGCs. This will not only improve the accuracy of annotations but also enable the discovery of larger and more complex BGCs that may have been missed using short-read methods. These expansions will provide a more comprehensive representation of the global chemical diversity of secondary metabolites and their relationship to environmental factors.

The data produced and made available in this work can also help in other areas of natural product research, and potentially become part of the BGC-Atlas in the future. One example application for the future is the bioactivity prediction from the BGCs identified in our database, using novel machine learning methods, such as NPF [28] and NPBdetect [29]. This can allow users to gain insights into bioactivity diversity correlations within a GCF and its associations with the environment. Beyond merely serving as a database, the BGC Atlas has the potential to become a foundational tool for hypothesis generation and experimental design in microbial secondary metabolite research, such as in the discovery of underexplored habitats with high biosynthetic potential, or in the study of how secondary metabolites drive microbial interactions at a global scale.

We encourage researchers to utilize the BGC Atlas to expand our understanding of microbial biosynthesis and its impact on ecological and evolutionary processes. The BGC Atlas is accessible at https://bgc-atlas.cs.unituebingen.de.

## 5 Data Availability

The resource is available and accessible at https://bgc-atlas.cs.uni-tuebingen.de. The resulting data is available on the website for download. The source code used to produce the data, and the web interface are available at https://github.com/ZiemertLab/bgc-atlas-analysis, and https://github.com/ZiemertLab/bgc-atlas-web, respectively, both with GNU General Public License v3.

## 6 Supplementary Data

Supplementary Data are available online.

## 7 ACKNOWLEDGEMENTS

The authors acknowledge the use of de.NBI cloud and the support by the High Performance and Cloud Computing Group at the Zentrum für Datenverarbeitung of the University of Tübingen and the Federal Ministry of Education and Research (BMBF) through grant no 031 A535A. We thank Interfaculty Institute for Biomedical Informatics (IBMI) at the University of Tübingen for providing the computational resources essential for this study. We also thank the National Supercomputing Mission (NSM) for providing computing resources of PARAM Smriti at NABI, Mohali, which is implemented by C-DAC and supported by the Ministry of Electronics and Information Technology (MeitY), Department of Biotechnology (DBT) and Department of Science and Technology (DST), Government of India. Additionally, we thank the Deutsche Forschungsgemeinschaft (DFG, German Research Foundation) under Germany’s Excellence Strategy—EXC 2124—390838134 for structural support. The icons that we use in the graphical abstract and the Figure 1 were downloaded from Flaticon.com, and the database schema for the Supplementary Figure 1 were drawn using draw.io.

## References

[1] Newman, D.J. and Cragg, G.M. (2016) Natural products as sources of new drugs from 1981 to 2014. Journal of Natural Products, 79(3), 629–661.

[2] Tyc, O., Song, C., Dickschat, J. S., Vos, M., and Garbeva, P. (2017) The ecological role of volatile and soluble secondary metabolites produced by soil bacteria. Trends in Microbiology, 25(4), 280–292.

[3] Salwan, R., and Sharma, V. (2020) Molecular and biotechnological aspects of secondary metabolites in actinobacteria. Microbiological Research, 231, 126374.

[4] Beppu, T. (1992) Secondary metabolites as chemical signals for cellular differentiation. Gene, 115(1-2), 159–165.

[5] Martin, J. F., and Liras, P. (1989) Organization and expression of genes involved in the biosynthesis of antibiotics and other secondary metabolites. Annual Review of Microbiology, 43(1), 173–206.

[6] Mungan, M. D., Alanjary, M., Blin, K., Weber, T., Medema, M. H., and Ziemert, N. (2020). ARTS 2.0: feature updates and expansion of the Antibiotic Resistant Target Seeker for comparative genome mining. Nucleic Acids Research, 48(W1), W546–W552.

[7] Yilmaz, T. M., Mungan, M. D., Berasategui, A., and Ziemert, N. (2023). FunARTS, the Fungal bioActive compound Resistant Target Seeker, an exploration engine for target-directed genome mining in fungi. Nucleic Acids Research, 51(W1), W191–W197.

[8] Streit, W. R., and Schmitz, R. A. (2004). Metagenomics–the key to the uncultured microbes. Current Opinion in Microbiology, 7(5), 492–498.

[9] Covington, B. C., Xu, F., and Seyedsayamdost, M. R. (2021). A natural product chemist’s guide to unlocking silent biosynthetic gene clusters. Annual Review of Biochemistry, 90(1), 763–788.

[10] Brady, S. F. (2007). Construction of soil environmental DNA cosmid libraries and screening for clones that produce biologically active small molecules. Nature Protocols, 2(5), 1297–1305.

[11] Geers, A. U., Buijs, Y., Strube, M. L., Gram, L., and Bentzon-Tilia, M. (2022) The natural product biosynthesis potential of the microbiomes of Earth – Bioprospecting for novel anti-microbial agents in the meta-omics era. Computational and Structural Biotechnology Journal, 20, 343–352.

[12] Mantri, S.S., Negri, T., Sales-Ortells, H., Angelov, A., Peter, S., Neidhardt, H., Oelmann, Y. and Ziemert, N. (2021) Metagenomic sequencing of multiple soil horizons and sites in close vicinity revealed novel secondary metabolite diversity. mSystems, 6(5), 10–1128.

[13] Paoli, L., Ruscheweyh, H.-J., Forneris, C. C., Hubrich, F., Kautsar, S., Bhushan, A., Lotti, A., Clayssen, Q., Salazar, G., Milanese, A., et. al. (2022) Biosynthetic potential of the global ocean microbiome. Nature, 607(7917), 111–118.

[14] Donia, M. S., Cimermancic, P., Schulze, C. J., Wieland Brown, L. C., Martin, J., Mitreva, M., Clardy, J., Linington, R. G., and Fischbach, M. A.(2014) A systematic analysis of biosynthetic gene clusters in the human microbiome reveals a common family of antibiotics. Cell, 158(6), 1402–1414.

[15] Gavriilidou, A., Kautsar, S. A., Zaburannyi, N., Krug, D., Müller, R., Medema, M. H., and Ziemert, N. (2022) Compendium of specialized metabolite biosynthetic diversity encoded in bacterial genomes. Nature Microbiology, 7(5), 726–735.

[16] Hover, B. M., Kim, S.-H., Katz, M., Charlop-Powers, Z., Owen, J. G., Ternei, M. A., Maniko, J., Estrela, A. B., Molina, H., Park, S., et. al. (2018) Culture-independent discovery of the malacidins as calcium-dependent antibiotics with activity against multidrug-resistant Gram-positive pathogens. Nature Microbiology, 3(4), 415–422.

[17] Feng, Z., Kallifidas, D., and Brady, S. F. (2011) Functional analysis of environmental DNA-derived type II polyketide synthases reveals structurally diverse secondary metabolites. Proceedings of the National Academy of Sciences, 108(31), 12629–12634.

[18] Hirsch, P., Tagirdzhanov, A., Kushnareva, A., Olkhovskii, I., Graf, S., Schmartz, G.P., Hegemann, J.D., Bozhüyük, K.A.J., Müller, R., Keller, A., Gurevich, A. (2024) ABC-HuMi: the Atlas of biosynthetic gene clusters in the human microbiome. Nucleic Acids Research, 52(D1), D579–D585.

[19] Blin, K., Shaw, S., Medema, M. H., and Weber, T. (2024) The antiSMASH database version 4: additional genomes and BGCs, new sequence-based searches and more. Nucleic Acids Research, 52(D1), D586–D589.

[20] Kautsar, S. A., Blin, K., Shaw, S., Weber, T., and Medema, M. H. (2021) BiG-FAM: the biosynthetic gene cluster families database. Nucleic Acids Research, 49(D1), D490–D497.

[21] Zou, H., Sun, T., Jin, B., and Wang, S. (2023). sBGC-hm: an atlas of secondary metabolite biosynthetic gene clusters from the human gut microbiome. Bioinformatics, 39(3), btad131.

[22] Richardson, L., Allen, B., Baldi, G., Beracochea, M., Bileschi, M.L., Burdett, T., Burgin, J., Caballero-Pérez, J., Cochrane, G., Colwell, L.J., et. al. (2023) MGnify: the microbiome sequence data analysis resource in 2023. Nucleic Acids Research, 51(D1), D753–D759.

[23] Blin, K., Shaw, S., Kloosterman, A. M., Charlop-Powers, Z., Van Wezel, G. P., Medema, M. H., and Weber, T. (2021) antiSMASH 6.0: improving cluster detection and comparison capabilities. Nucleic Acids Research, 49(W1), W29–W35.

[24] Kautsar, S. A., van der Hooft, J. J., de Ridder, D., and Medema, M. H. (2021) BiG-SLiCE: A highly scalable tool maps the diversity of 1.2 million biosynthetic gene clusters. GigaScience, 10(1), giaa154.

[25] Terlouw B.R., Blin K., Navarro-Munoz J.C., Avalon N.E., Chevrette M.G., Egbert S., Lee S., Meijer D., Recchia M.J., Reitz Z.L. et. al. (2023) MIBiG 3.0: a community-driven effort to annotate experimentally validated biosynthetic gene clusters. Nucleic Acids Research, 51(D1), D603–D610.

[26] Chen, I.-M. A., Chu, K., Palaniappan, K., Ratner, A., Huang, J., Huntemann, M., Hajek, P., Ritter, S. J., Webb, C. Wu, et. al. (2023) The IMG/M data management and analysis system v. 7: content updates and new features. Nucleic Acids Research, 51(D1), D723–D732.

[27] Chikhi, R., Raffestin, B., Korobeynikov, A., Edgar, R. C., and Babaian, A. (2024) Logan: Planetary-Scale Genome Assembly Surveys Life’s Diversity. bioRxiv, 2024–07.

[28] Walker, A. S., and Clardy, J. (2021) A machine learning bioinformatics method to predict biological activity from biosynthetic gene clusters. Journal of Chemical Information and Modeling, 61(6), 2560–2571.

[29] Goyat, H., Singh, D., Paliyal, S., and Mantri, S. S. (2024) Predicting biological activity from biosynthetic gene clusters using neural networks. bioRxiv, 2024–06.

