## Supplementary Data for "BGC Atlas: A Web Resource for Exploring the Global Chemical Diversity Encoded in Bacterial Genomes"

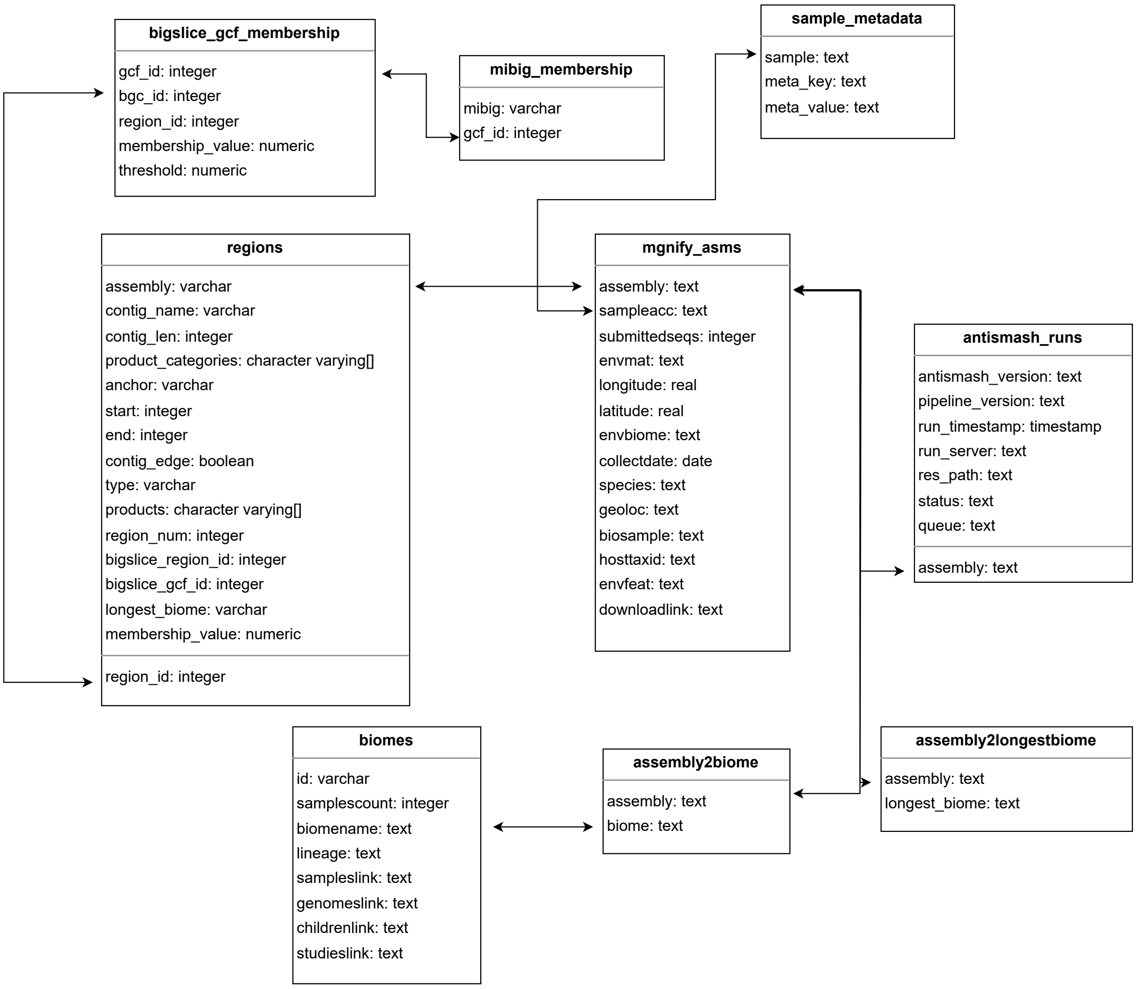


Figure 1: Database Schema of BGC Atlas. This diagram illustrates the structure of the database used in the BGC Atlas, including key tables and relationships between them. The schema encompasses tables for MGnify assemblies, antiSMASH runs, biosynthetic gene cluster (BGC) memberships of gene cluster families (GCF), biome associations, and metagenomic assembly metadata. Each table is linked through unique identifiers.

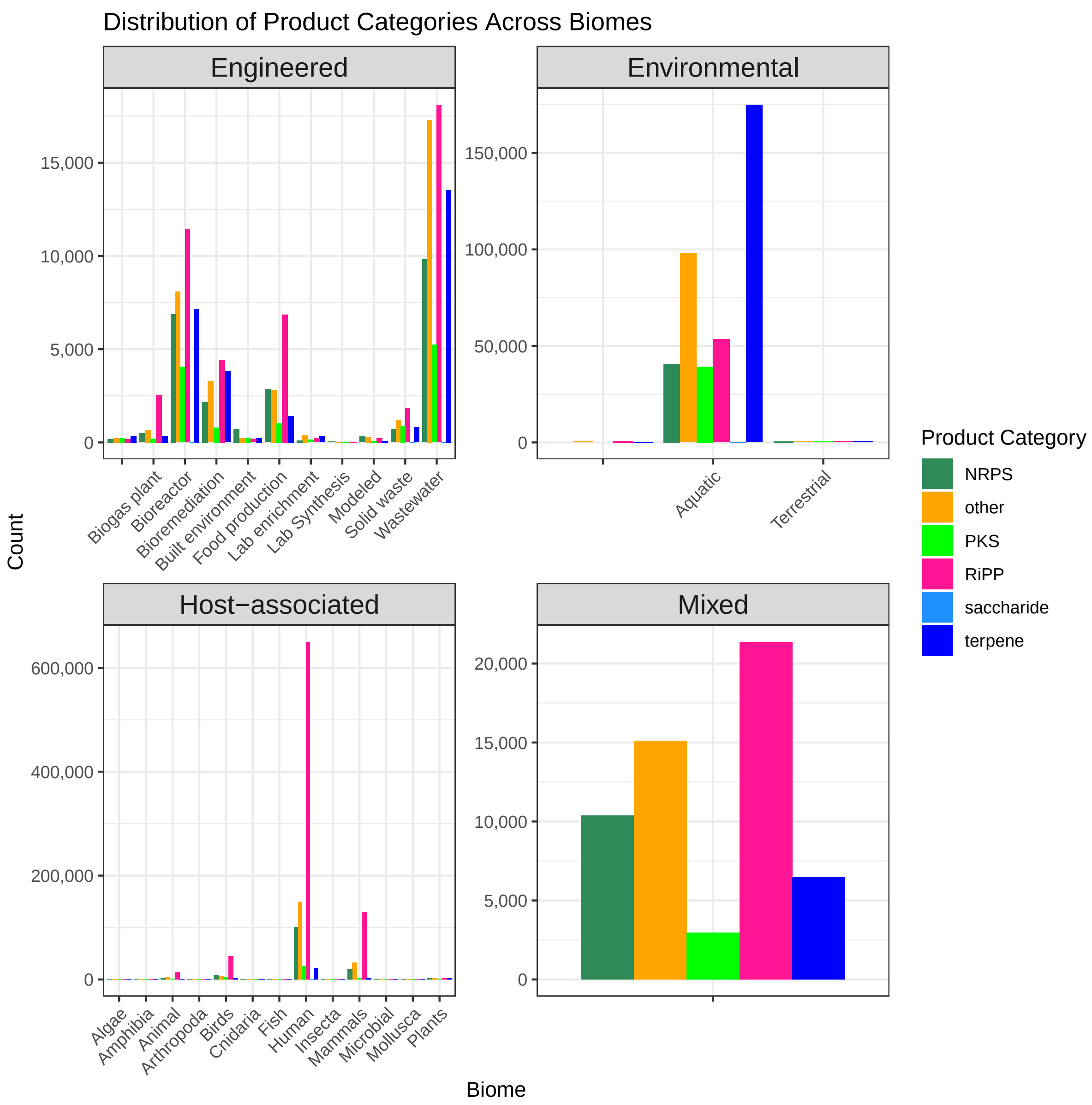


Figure 2: Distribution of product categories across biomes. The figure displays the distribution of various product categories, including nonribosomal peptides (NRPS), polyketides (PKS), ribosomally synthesized and post-translationally modified peptides (RiPPs), saccharides, terpenes, and others, across different biome types. The biomes are grouped into four major categories: host-associated, mixed, engineered, and environmental, which are the first level of the biome hierarchy. The x-axis for each for first-level category shows the second-level categories of them. The data highlights the variability in product category prevalence across distinct biomes, with certain categories being more abundant in specific environmental contexts.
